# Presaccadic attention sharpens visual acuity

**DOI:** 10.1101/2022.11.02.514935

**Authors:** Yuna Kwak, Nina M. Hanning, Marisa Carrasco

**Affiliations:** Department of Psychology, NYU; Neural Science, NYU; Department of Psychology, Humboldt-Universität zu Berlin, Berlin, Germany

## Abstract

Visual perception is limited by spatial resolution, which declines with eccentricity and differs around polar angle locations. To compensate for poor peripheral resolution, we make rapid eye movements –saccades– to bring peripheral objects into high-acuity foveal vision. Already before saccade onset, visual attention shifts to the saccade target location and prioritizes visual processing. This *presaccadic shift of attention* improves performance in many visual tasks, but whether it changes resolution is unknown. Here, we investigated whether presaccadic attention sharpens peripheral spatial resolution; and if so, whether such effect interacts with polar angle locations. We measured acuity thresholds in an orientation discrimination task during fixation and saccade preparation around the visual field. The results revealed that presaccadic attention sharpens acuity, which can facilitate a smooth transition from peripheral to foveal representation. This acuity enhancement is similar across the four cardinal locations; thus, presaccadic attention does not change polar angle differences in resolution.

## Introduction

The human visual system processes the world with a great degree of spatial non-uniformity. Our spatial resolution, the ability to discriminate fine spatial patterns, is highest at the fovea and decreases with eccentricity (for reviews, ^1,2^). Importantly, visual perception not only varies with eccentricity but also as a function of polar angle – known as ‘performance fields’: at iso-eccentric locations, visual performance is better at the horizontal than the vertical meridian (HVA: horizontal-vertical anisotropy) and is better at the lower vertical than the upper vertical meridian (VMA: vertical meridian asymmetry). Such non-uniformities across eccentricity and polar angle have been reported for various visual tasks and dimensions ^3–10^, including spatial resolution ^11–14^.

Visual perception is limited by spatial resolution. One of the ways that we can compensate for this limit is by making rapid movements –saccades– to position behaviorally relevant stimuli at the fovea, where we have the highest resolution. As spatial resolution decreases with eccentricity, a blurry representation in the periphery (presaccade) will appear in high detail once brought into the fovea by a saccade (post-saccade). Interestingly, visual attention shifts to the saccade target already before the eyes move, and selectively prioritizes visual processing at the upcoming fixation location ^15,16^. The presaccadic shift of attention has been reported to enhance visual performance ^17,18^, perceived contrast ^19^ and contrast sensitivity ^20,21^, and reshapes sensory tuning of basic visual features ^22–24^. These presaccadic modulations can facilitate perceptual stability-a smooth transition from presaccadic (peripheral) to post-saccadic (foveal) information– by making the peripheral representation become more fovea-like in anticipation of eye movements ^22–26^.

Given the prevalence of presaccadic attention in selective processing of visual information and that spatial resolution changes drastically from foveal to peripheral visual field, here we investigate whether presaccadic attention changes spatial resolution. Deploying *covert* endogenous (voluntary) or exogenous (involuntary) spatial attention (i.e., without concurrent eye movements) increases performance in spatial resolution tasks ^12,27–31^. Montagna and colleagues (2009), for example, asked to report the location of a gap in a Landolt-square. They found that both endogenous and exogenous attention led to smaller gap-size thresholds at the attended and larger at the unattended locations compared to the neutral condition, indicating a trade-off in spatial resolution with both types of attention. Given that both covert attention and presaccadic attention improve performance, one might hypothesize that presaccadic attention also enhances spatial resolution. However, the perceptual effects of presaccadic attention and covert attention are dissociable in various ways (for review, ^32^). They rely on differential neural computations and recruit separable neural populations ^21,33–35^. Presaccadic and covert attention also differentially modulate the representation of basic visual features, such as orientation and spatial frequency ^22–24^. Given these differences, it remains an open question whether presaccadic attention modulates spatial resolution.

The aim of the present study was two-fold. First, we investigated whether presaccadic attention increases spatial resolution using a visual acuity task. Second, we examined whether this effect interacts with the known ‘performance fields’asymmetries across polar angle – horizontal-vertical anisotropy (HVA) and vertical meridian asymmetry (VMA). These asymmetries are ubiquitous: they emerge for different orientations, spatial frequencies, and eccentricities ^9–11,36^, when a target is presented by itself or among distractors ^9,36,37^; monocularly and binocularly ^9,11^; and for different head tilts ^38^, and are preserved during visual working memory ^13,39^. Moreover, they are resilient: deploying covert attention either endogenously or exogenously benefits performance similarly across polar angle locations, thus preserving the shape of the performance fields ^9,36,37,40,41^. Likewise, presaccadic attention may also preserve polar angle asymmetries in spatial resolution. Given that presaccadic and covert attention effects are dissociable ^32^, it is also possible that presaccadic attention alleviates these asymmetries by enhancing performance more at locations where perception could benefit most. Alternatively, based on a recent finding that presaccadic attention even exacerbates performance field asymmetries in contrast sensitivity ^20^, presaccadic attention may exacerbate them in spatial resolution too.

Here, to investigate whether and how presaccadic attention modulates spatial resolution around the visual field, we compared visual acuity thresholds during fixation and saccade preparation for each of the four cardinal locations. We found that presaccadic attention increases acuity thresholds already before saccade onset, consistent with its role in narrowing the gap between peripheral and foveal representations. The acuity benefit was similar around the visual field, indicating that performance field asymmetries in acuity are resilient and cannot be allayed by presaccadic attention.

## Results

We measured visual acuity at the left, right, upper, and lower visual field (**Figure 1A**) when participants prepared to look toward the test stimulus (*valid*), maintained fixation (*baseline*), or prepared to look away from the test stimulus (*invalid*). The task was to judge the orientation (CW or CCW) of a full contrast Gabor stimulus, of which we adjusted the spatial frequency using adaptive staircasing procedures, to determine the spatial frequency leading to 75% accuracy (acuity threshold) – separately for each combination of location and eye movement condition (**Figure 1B**; see Materials and Methods – Titration procedures).

**Figure 1.**
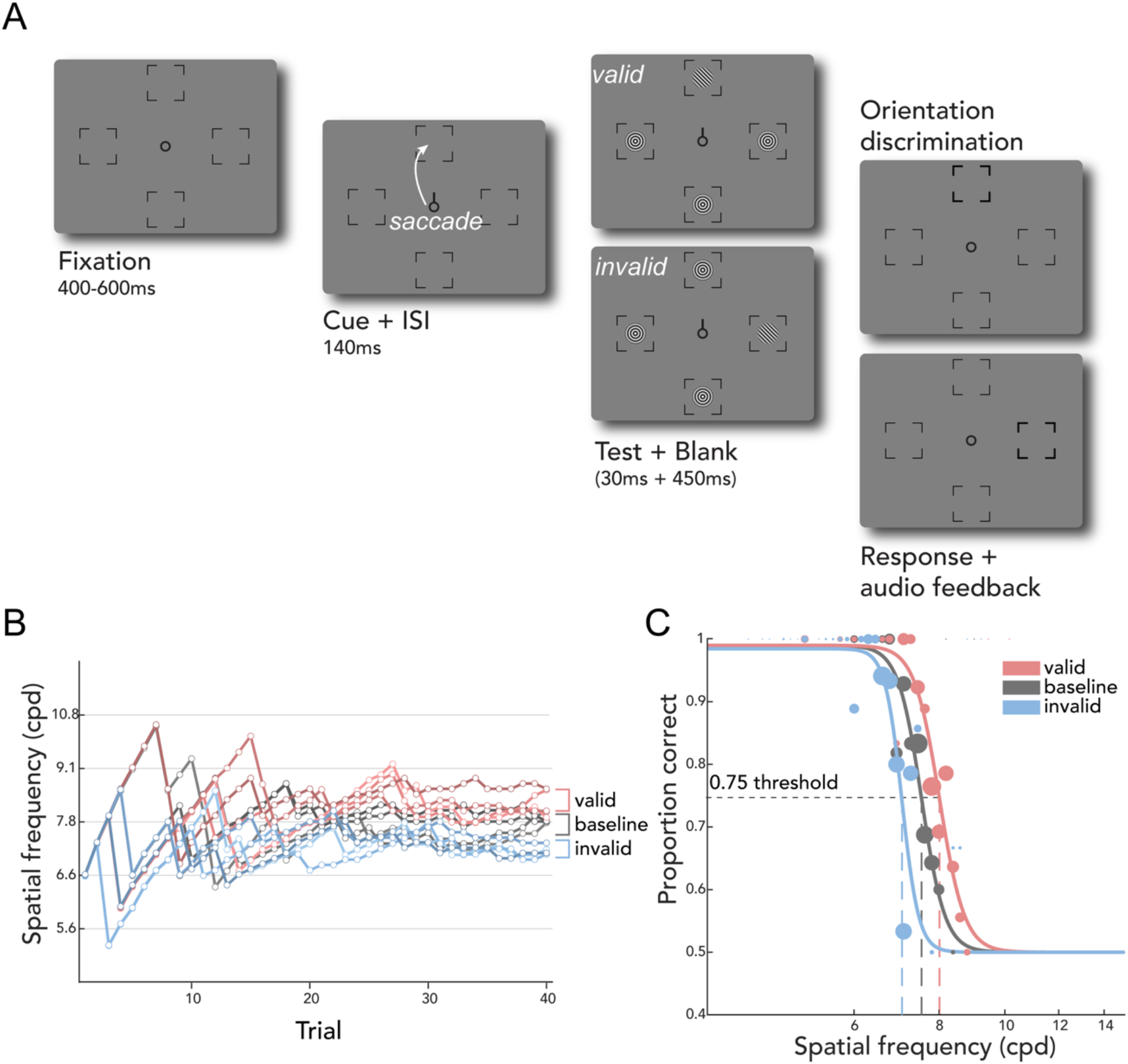
Experimental design **A)** Trial sequence for eye movement blocks. After a fixation period, a central saccade cue indicated the saccade target. Participants were instructed to make a saccade to the center of the corresponding placeholder as soon as the cue appeared. Just before saccade onset, one test Gabor stimulus and three radial frequency pattern distractors were presented briefly. The test Gabor could appear at the cued (*valid*) or at one of the un-cued (*invalid*) locations. After a delay, a boldened placeholder indicated the location at which the test Gabor was presented, and participants judged its orientation (CW or CCW). The spatial frequency of the test and the matching distractors was varied based on Bayesian adaptive procedures aimed at finding the 75% acuity threshold (Figure 1B). In the fixation blocks (*baseline;* not illustrated here), the trial sequence was identical except that the cue pointed to all four locations, and participants were instructed to fixate at the center throughout the entire block. **B)** Adaptive staircases for a representative participant for one location. For each location, we ran four independent staircases per condition (*valid, baseline*, and *invalid*). The estimated 75% thresholds for the staircases were averaged to derive the final acuity thresholds. **C)** The responses from the adaptive staircases were used to fit a psychometric function for each condition-location combination to derive 75% threshold and slope parameters. The size of each colored dot corresponds to the number of trials in each bin.

We first evaluated visual acuity during fixation. The current data reproduced robust performance field asymmetries in visual acuity ^11^. **Figure 2A** shows the group average normalized acuity thresholds, computed by normalizing each participant’s acuity threshold at a location by their average of thresholds at all locations. During fixation, participants had higher visual acuity at the horizontal than at the vertical meridian (mean difference in cpd ±SEM: 2.155 ±0.199), consistent with the horizontal-vertical anisotropy (HVA). In line with the vertical meridian asymmetry (VMA), visual acuity was also higher for the lower than the upper visual field (mean difference in cpd ±SEM: 0.790 ±0.186).

**Figure 2.**
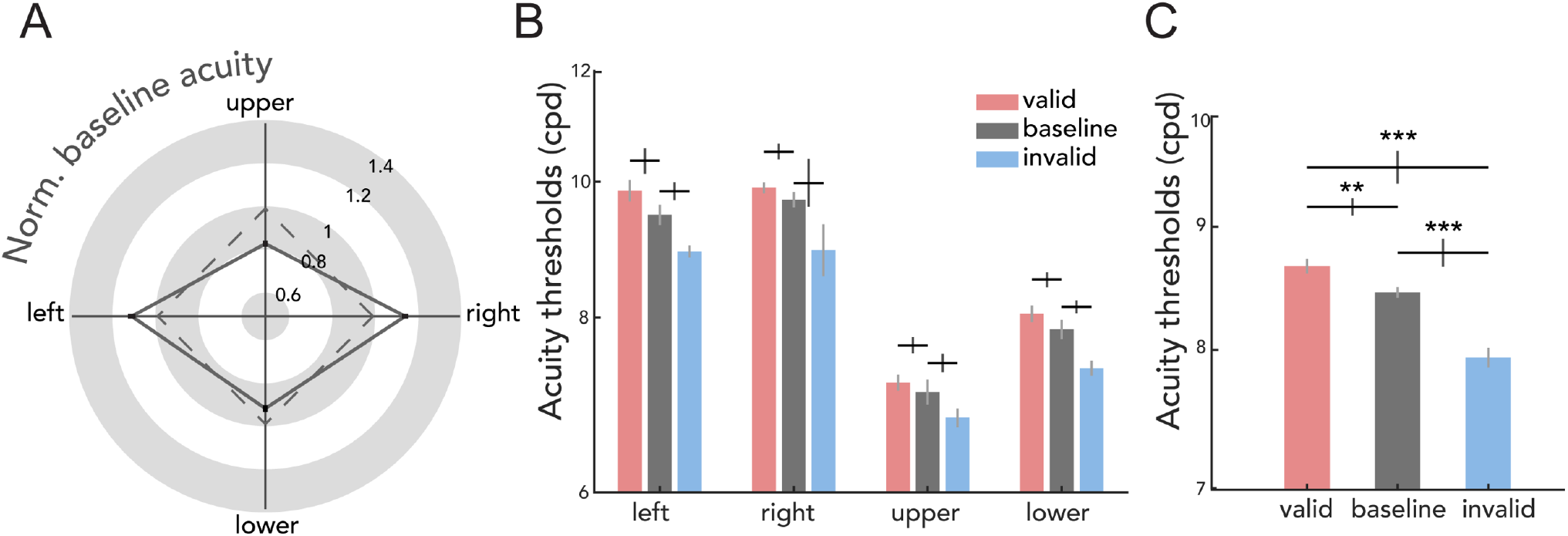
Main results. **A)** Group averaged acuity thresholds in the *baseline* condition for each location normalized by participants’ averaged *baseline* threshold across (solid line). The black lines at each location depict ±1 SEM. The dashed line illustrates a scenario in which there would be no asymmetries. Baseline visual acuity is better at the horizontal than the vertical meridian, and at the lower visual field than the upper visual field. **B)** Acuity thresholds for each condition (*valid, baseline, invalid*) and location. Each condition demonstrates typical performance field asymmetries. **C)** Acuity thresholds for each condition averaged across locations. Presaccadic acuity benefits at the saccade target (*valid*) and costs at non-target locations (*invalid*) compared to fixation (*baseline*). Y-axis is plotted in log-scale. Gray error bars in B & C depict ±1 SEM (Cousineau corrected), black error bars depict ±1 SEM of the difference between the compared conditions. *\*\*p*< 0.01, \*\*\**p*< 0.001, FDR corrected.

Next, we evaluated visual acuity benefits and costs during saccade preparation relative to fixation. **Figure 2B** shows the visual acuity thresholds in units of cpd for each condition-location combination. To test for the effect of presaccadic attention around the visual field, we conducted a 3 (*valid, invalid, baseline*) by 4 (left, right, upper, lower) repeated-measures ANOVA, which yielded main effects of condition (*F*(2,22) = 26.969, *p* < 0.001, BF10 > 100) and location (*F*(3,33) = 90.556, *p* < 0.001, BF10 > 100), but no significant interaction (*F*(6,66) = 0.810, *p* = 0.587, BF10 = 0.057). This indicates that acuity was independently modulated by saccade preparation and polar angle location. To further explore the effect of presaccadic attention independent of polar angle, we combined the data across the 4 locations (**Figure 2C**). Compared to fixation (*baseline*), visual acuity was higher for the *valid* and lower for the *invalid* condition (*valid-baseline t*(11) = 2.794, *p* = 0.005, BF10 = 3.778; *baseline-invalid t*(11) = 5.174, *p* < 0.001, BF10 > 100; *valid-invalid t*(11) = 5.742, *p* < 0.001, BF10 > 100). This pattern was consistent across individuals and across polar angle locations (**Figure 3**). The respective benefits and costs in visual acuity while preparing saccades toward and away from the test stimulus demonstrate that presaccadic attention enhances spatial resolution, and these effects are uniform across the visual field.

**Figure 3.**
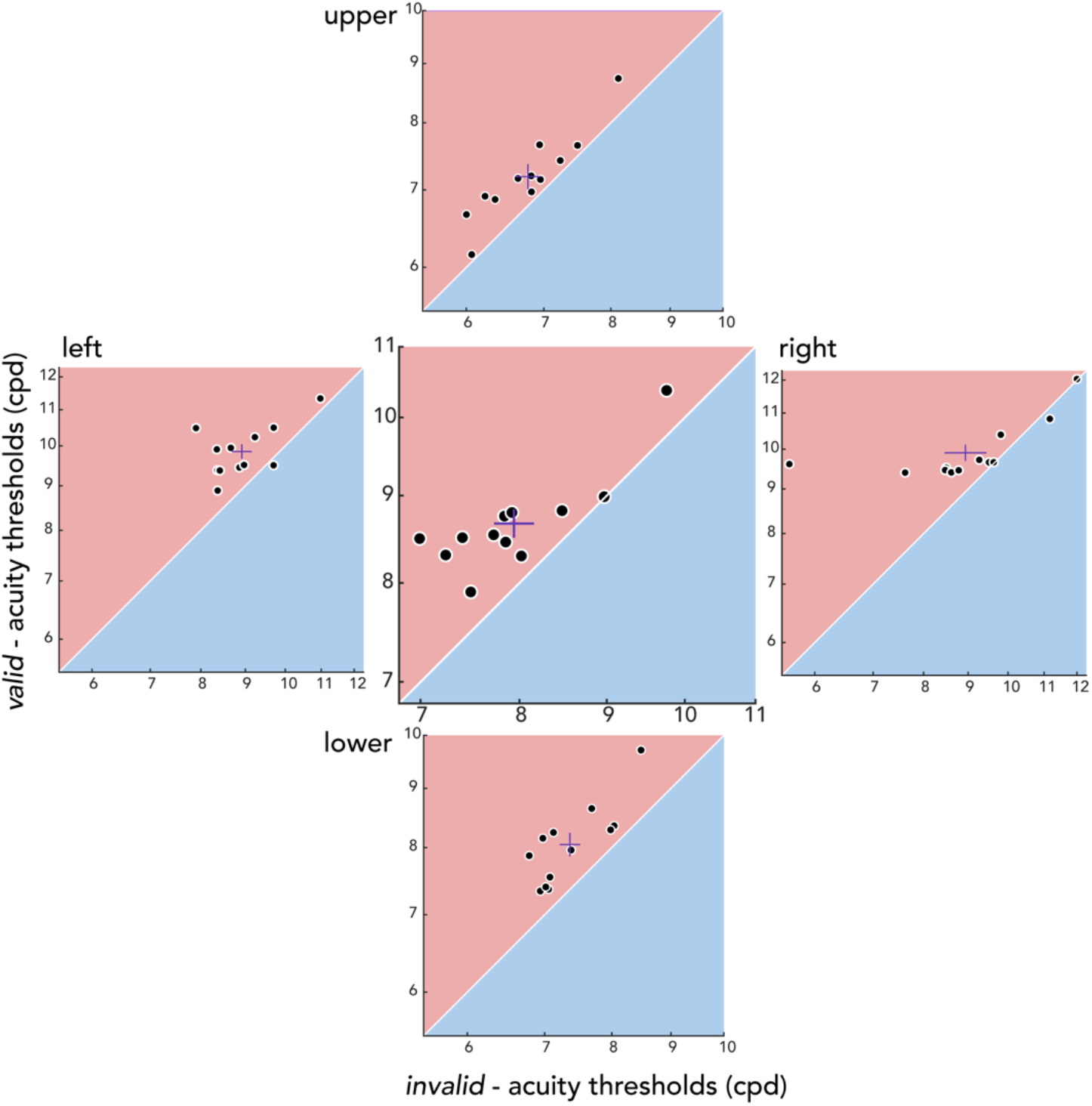
Individual participant’s presaccadic acuity. Benefits and costs for each location (outer plots) and averaged across locations (center plot). Thresholds for the *valid* condition (y-axis) are plotted against the *invalid* condition (x-axis), and both axes are in log scale. Red lines depict mean ±1 SEM. Note that the range plotted differs between locations, to account for visual field asymmetries.

Because the adaptive staircase procedures do not allow for a detailed evaluation of eye movement parameters ‘online’ during the ongoing experiment, we also performed an offline eye movement analysis after the experiment using a more precise saccade detection algorithm ^42^ (see Materials and Methods – Eye movement data analysis). To examine whether the staircase acuity thresholds derived online would hold after excluding trials (7.4% of all trials) in which the test stimulus was not presented within the last 150ms prior to saccade onset according to the offline algorithm, we fit a psychometric function for each location-condition combination using trials from all 4 staircases that survived the offline algorithm (**Figure 1B-C**). Importantly, acuity thresholds results were very similar regardless of whether the trials filtered out from the offline analysis were included (adaptive staircase; **Figure 2B**) or excluded (psychometric function fitting).

The online staircase-derived thresholds including all trials (**Figures 1B, 2B**) were correlated with the threshold estimates derived from psychometric function fitting after excluding additional trials based on the abovementioned offline criteria (**Figure 1C**; *r* = 0.964, *p* < 0.001, BF10 > 100). Moreover, as with the thresholds derived from the adaptive staircase procedures, a 2-way repeated measures ANOVA on the fitted thresholds showed main effects of condition (*F*(2,22) = 26.79, *p* < 0.001, BF10 > 100) and location (*F*(3,33) = 84.66, *p* < 0.001, BF10 > 100), but no interaction effect (*F*(6,66) = 0.867, *p* = 0.524, BF10 = 0.054). In addition, we evaluated the slope parameters extracted from the fitted acuity functions. There were neither main effects (condition: *F*(2,22) = 1.586, *p* = 0.227, BF10 = 0.204; location: *F*(3,33) = 1.401, *p* = 0.260, BF10 = 0.202) nor an interaction effect (*F*(6,66) = 1.821, *p* = 0.108, BF10 = 0.928). Therefore, consistent with previous work ^11^, the slope was constant across visual field locations during fixation. Moreover, it was unaffected by presaccadic attention.

We also evaluated eye movement parameters for different saccade directions (**Figure 4**). A repeated-measures ANOVA showed a significant main effect of saccade direction on saccade latency (*F*(3,33) = 13.531, *p* < 0.001, BF10 > 100). Downward saccades were slower than saccades toward other locations (leftward *t*(11) = 4.945, *p* < 0.001, BF10 = 80.729; rightward *t*(11) = 4.326, *p* = 0.002, BF10 = 34.292; upward *t*(11) = 6.170, *p* < 0.001, BF10 > 100), consistent with previous research ^20,43–45^. There was also a significant main effect of saccade direction on saccade amplitude (*F*(3,33) = 3.501, *p* = 0.025, BF10 = 3.527). Rightward saccades had higher amplitudes than leftward (*t*(11) = 3.775, *p* = 0.012, BF10 = 15.618), upward (*t*(11) = 2.407, *p* = 0.027, BF10 = 2.192), and downward saccades (*t*(11) = 2.403, *p* = 0.036, BF10 = 2.180), consistent with amplitude asymmetries reported previously ^20,46^. Previous work however has shown that presaccadic attention shifts to the intended saccade target, independent of the ultimately executed saccade program ^15,47^; thus the quality of presaccadic attention is likely to be unaffected latency and amplitude differences across locations. Moreover, there was no significant difference in landing precision (mean of the Euclidean distance between saccade endpoints and saccade target center) across the four saccade directions (*F*(3,33) = 0.500, *p* = 0.681, BF10 = 0.177), also in line with a previous study ^20^. Therefore, even with latency and amplitude varying with saccade direction in a manner consistent with previous work, the presaccadic modulation of acuity was consistent across all polar angle locations (**Figures 2B-C, 3**).

**Figure 4.**
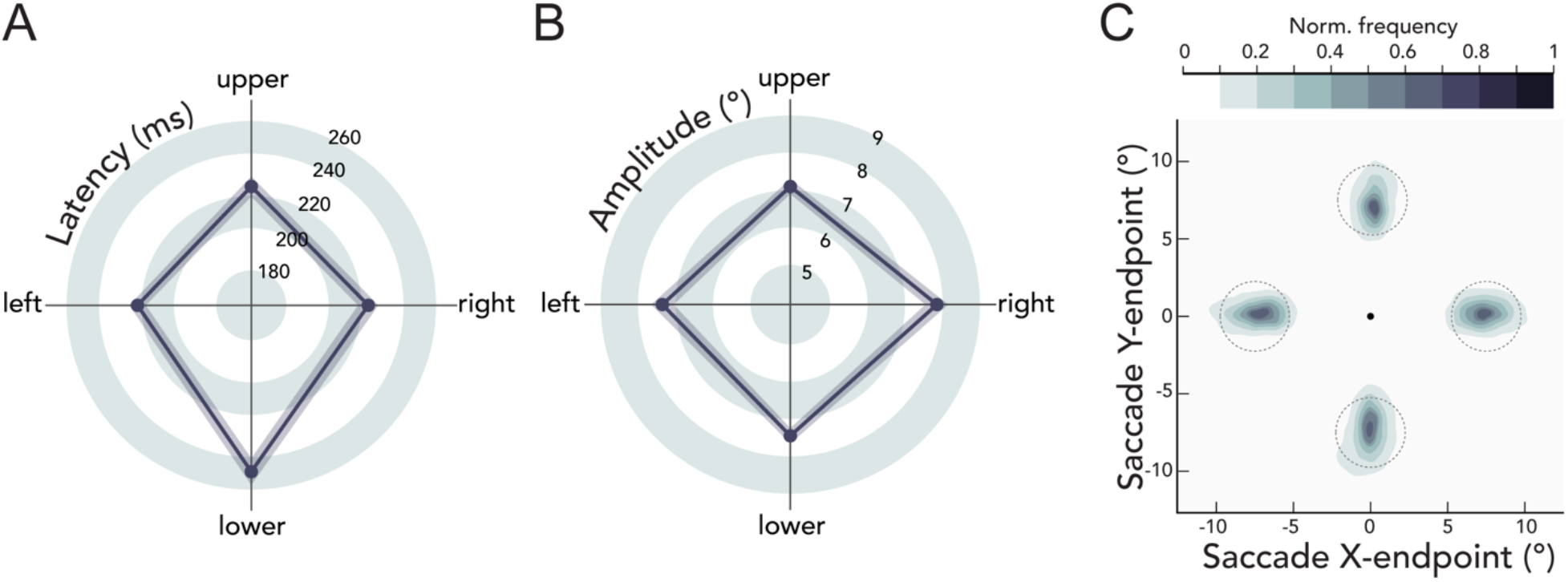
Eye movement parameters. Group average **A)** saccade latencies (ms) and **B)** saccade amplitudes (°) as a function of saccade direction. Shaded error areas indicate ±1 SEM. **C)** Normalized saccade endpoint frequency maps averaged across participants as a function of saccade direction. Dashed gray circles indicate the 2.25°saccade target region. The black circle in the center indicates the fixation location.

## Discussion

In the current study, we asked whether presaccadic attention enhances spatial resolution in a visual acuity task, and if so, whether presaccadic benefits and costs differ across polar angle locations. Before saccade onset, visual acuity was enhanced at the saccade target (presaccadic benefits) and reduced for non-target locations (presaccadic costs), compared to baseline acuity measured during fixation. This presaccadic attentional effect was uniform across the four cardinal locations, demonstrating that the acuity asymmetries around the visual field are resilient and cannot be overcome even with the typically robust effect of saccade preparation.

Directing covert spatial attention, without eye movements, increases performance in various spatial resolution tasks, e.g., using Landolt-squares, broken lines, vernier offsets or texture segmentation ^12,27–31^. Covert attention improving performance in resolution tasks has been attributed to enhanced quality of target representations ^48,49^, rather than a change in decisional criteria or efficient inhibition of non-relevant information ^29,50,51^. Notably, covert endogenous and exogenous attention differ in how they modulate spatial resolution (for review, see ^49^): whereas endogenous attention is a flexible mechanism that can increase but also decrease resolution depending on task demands ^52,53^, exogenous attention automatically enhances resolution, even if detrimental to the task ^14,31,54^.

Here, we found evidence that presaccadic attention –which is dissociable from covert attention– also enhances spatial resolution. Presaccadic effects on acuity were small but reliable: on average, benefits and costs combined across locations were 0.213 cpd (~2.5%) and 0.504 cpd (~5.9%), respectively. We cannot directly compare the magnitude of this effect to that of covert attention because enhanced resolution with covert attention has been measured with tasks in which spatial frequency is not the dependent measure. However, the magnitude of the presaccadic effects reported here are similar to those of exogenous ^55^ and endogenous ^56^ covert attention on *apparent* spatial frequency.

Increased visual acuity at the saccade target is consistent with previous reports of presaccadic attention selectively enhancing sensitivity to higher spatial frequencies ^22,57^. In particular, Li et al. (2016) showed that the peak of the spatial frequency tuning curve shifts toward higher frequencies during saccade preparation. Given that high spatial frequency filters directly relate to our ability to discriminate fine spatial details, enhanced sensitivity of these filters is consistent with increased visual acuity thresholds during saccade preparation. Moreover, presaccadic attention enhances processing of high spatial frequencies even if this results in impaired performance ^22^. Thus, the sharpened acuity observed in the current study is likely to occur automatically, regardless of the task at hand.

Presaccadic attention enhancing sensitivity to high spatial frequencies may relate to the contribution of neurons representing the foveal region –which are particularly sensitive to these spatial frequencies– in processing the peripheral target already before saccade onset in anticipation of the post-saccadic visual input. In fact, neurophysiological evidence shows that just before the onset of a saccadic eye movement, neurons in visual and parietal cortex tuned to the saccade target location increase their firing rate in a predictive manner ^58–61^, and receptive fields of neurons in visual and prefrontal cortex are shifted in location toward the saccade target ^62–64^.

Furthermore, a study testing a broad range of spatial frequencies revealed that presaccadic attention reshapes the spatial frequency sensitivity profile, such that the bandwidth becomes narrower ^57^: presaccadic benefits increased for spatial frequencies from 1.0 to 2.5 cpd and gradually decreased, but were still present at 5.5 cpd (the highest spatial frequency tested). Note that the spatial frequencies used in the present study are well above those tested in previous work ^22,23,57^. This is because our study was aimed at examining acuity per se –around the spatial frequency range which is closer to the natural bounds of spatial resolution at a given eccentricity ^65^. Moreover, whereas previous studies mentioned above titrated stimulus contrast to avoid ceiling or flooring in performance, stimuli in the current study were fixed at full contrast to minimize the effect of contrast sensitivity and to define performance only by the spatial frequency dimension. The combination of high spatial frequencies and high contrast enabled us to selectively target visual acuity by isolating the presaccadic effects on spatial resolution from its effect on contrast-sensitivity.

Another aim of the study was to investigate whether presaccadic attention modulates polar angle asymmetries in acuity. Our results demonstrate that presaccadic attention enhances acuity similarly across the visual field. As a consequence, even with the deployment of presaccadic attention, visual acuity remains considerably worse at the upper vertical meridian, where baseline performance is usually worst and thus could have benefitted more than other locations. Thus, the typical shape of the acuity asymmetries (**Figure 2A**) could not be compensated by preparing saccades, demonstrating once again the robustness of the asymmetries. Likewise, neither endogenous nor exogenous attention can mitigate performance field asymmetries in texture segmentation ^14^, contrast sensitivity ^36,37^, and acuity tasks ^12^, for neurotypical participants as well as for those with amblyopia ^40^ or attention-deficit hyperactivity disorder ^41^. Such impervious constraints in visual performance may be explained by anatomical restraints in the cortex: there is substantially more cortical surface area dedicated to processing the horizontal than the vertical meridian, and the lower than the upper vertical meridian ^66,67^. Our results suggest that the asymmetries in neural resources, which may underlie the behavioral asymmetries, cannot be compensated for by presaccadic attention.

A recent study investigated presaccadic modulation of contrast sensitivity, using similar adaptive staircase procedures and experimental parameters as in the current study ^20^. As with covert spatial attention ^9,36,37,40,41^, presaccadic attention did not alleviate polar angle asymmetries in contrast sensitivity thresholds. Rather, the asymmetry was exacerbated: there were no presaccadic benefits before upward saccades, i.e., at the upper vertical meridian where contrast sensitivity is worst. This pattern is different from those reported with covert attention and also from the presaccadic acuity benefits reported here for all locations, including the upper visual field. However, these seemingly contradictory effects of presaccadic attention on contrast sensitivity and acuity may be explained by the fact that the two studies are targeting different points on the contrast sensitivity function (CSF). The CSF is characterized by a joint manipulation of contrast and spatial frequency, but both studies examined the presaccadic modulation of only one of the two, while fixing the other. Theoretically, visual acuity measured in the present work corresponds to the cut-off point of the CSF –the spatial frequency at which full contrast is needed for 75% orientation discriminability– and thus we fixed contrast while varying spatial frequency. Hanning and colleagues (2022) however tested relatively closer to the peak of the CSF, by fixing spatial frequency and varying contrast. Perhaps the manner with which the CSF is reshaped with presaccadic attention differs depending on polar angle location. Further research is needed for a full picture on how acuity, contrast sensitivity, and performance fields interact –particularly in relation to presaccadic attention.

In conclusion, the present findings reveal that presaccadic attention sharpens visual acuity at the saccade target already before it is brought into high-resolution foveal vision. By narrowing the difference between foveal and peripheral acuity, this shift facilitates a smooth transition between peripheral and foveal representations. These results parallel the remarkable stability with which visual percepts are maintained across rapid changes of retinal images during saccadic eye movements. The boost in acuity was uniform at iso-eccentric locations around the visual field, thus preserving rather than changing the shape of visual field asymmetries, despite the typically robust effect of presaccadic attention.

### Limitations of the study

The present study cannot speak to why and how presaccadic attention preserves performance field asymmetries by boosting acuity similarly around the visual field, whereas it exacerbates contrast sensitivity asymmetries by enhancing sensitivity around polar angle locations but the upper visual field ^20^. In addition, it would be interesting to evaluate the time course of presaccadic benefits-i.e., enhancement in performance as the test stimulus appears closer in time to saccade onset-around the visual field. The current experimental design leveraging adaptive staircase procedures was not optimized for distributing trials across time points before saccade onset. Whether presaccadic benefits for acuity evolve over time remains a question for future research.

## Materials and Methods

### Participants

Twelve observers (7 females, including authors YK and NH, ages 20-32) with normal or corrected-to-normal vision participated in the experiment. All participants (except for the two authors) were naïve to the purpose of the experiment and were paid $12 per hour. The Institutional Review Board at New York University approved the experimental procedures, and all participants provided written consent.

### Setup

Participants sat in a dimly lit room with their head stabilized by a chin and forehead rest and viewed the stimuli at 79 cm distance. All stimuli were generated and presented using MATLAB (MathWorks, Natick, MA, USA) and the Psychophysics Toolbox ^68,69^ on a gamma-linearized 20-inch ViewSonic G220fb CRT screen (Brea, CA, USA) with a spatial resolution of 1,280 by 960 pixels and a refresh rate of 100 Hz. Gaze position was recorded using an EyeLink 1000 Desktop Mount eye tracker (SR Research, Osgoode, Ontario, Canada) at a sampling rate of 1 kHz.

### Experimental design

**Figure 1A** shows the experimental design used to examine whether presaccadic attention modulates spatial resolution across the visual field. We assessed spatial resolution by measuring visual acuity (or spatial frequency) thresholds resulting in 75% accuracy in an orientation discrimination task. There were two eye movement conditions (*valid* and *invalid*) and a fixation condition (*baseline*). The two eye movement conditions were randomly intermixed within each block, whereas the fixation condition was run in separate blocks.

Each trial started with a fixation circle (radius 0.175°) on a gray background (~26 cd/m^2^) with a duration randomly jittered between 400ms and 600ms. There were four placeholders indicating the locations of the upcoming stimuli, 7.5°left, right, above, and below fixation. Each placeholder was composed of four corners (black lines, length 0.2°) delimiting a virtual square (distance between corners 5.6°). The trial began once a 300-ms stable fixation (eye coordinates within 2.25°radius virtual circle centered on fixation) was detected.

In eye movement blocks (*valid* and *invalid*), after the fixation period, a central direction cue (black line, length 0.4°) pointed to one of the four cardinal placeholders, cueing the saccade target. Participants were instructed to move their eyes to the center of the cued location as fast and precisely as possible. 140ms after cue onset (i.e., before saccade onset), a test Gabor grating (100% contrast, tilted ±45 relative to vertical, random phase, delimited by a raised cosine envelope with radius 2°) appeared for 30ms either at the cued saccade target location (*valid*,50% of trials), or at one of the other three non-target locations randomly chosen on each trial (*invalid*, 50% of trials). This timing ensured that the stimulus offset would be within a reasonable time window from saccade onset. Trials in which saccades were initiated before stimulus offset were excluded and repeated at the end of each block. The spatial frequency of the test Gabor was titrated using an adaptive psychometric staircase procedure (**Figure 1B**, see Materials and Methods-Titration procedures). To prevent biasing eye movements to a single sudden-onset stimulus, radial frequency patterns (i.e., containing no orientation information) matching the spatial frequency, contrast, and phase of the Gabor, appeared at the other three locations where the test Gabor did not appear. 450ms after stimuli offset, the test location was highlighted by increasing the width and length of the corresponding placeholders. Participants performed an un-speeded, orientation discrimination task (left arrow key for −45°, right arrow key for +45°), and received auditory feedback as to whether they were correct or incorrect.

Stimulus parameters and timing for the fixation blocks (*baseline*) were identical to the eye movement blocks with one exception: instead of one saccade target cue, four black direction cue lines pointed to all locations. Participants kept fixation throughout the entire trial sequence.

The experiment started with a fixation block, to measure an initial spatial frequency threshold for each location, which was used to determine the starting point of the staircases for the following blocks (see Materials and Methods-Titration procedures). Afterwards, participants completed four fixation blocks (160 trials per block) and four eye movement blocks (320 trials per block) that were randomly interleaved. Gaze position was monitored online to ensure correct fixation and precise eye movements. In fixation blocks and the fixation period of eye movement blocks, trials in which gaze deviated from 2.25°radius of fixation were aborted and repeated at the end of each block. In eye movement blocks, we additionally repeated trials with too short (< 150ms) or long (> 350ms) saccade latencies or missing / incorrect eye movements (saccade not landing within 2.25°from the center of the cued location). We collected a total of 1920 trials per participant-1280 eye movement trials (640 *valid*, 640 invalid) and 640 fixation trials.

### Titration procedures

We titrated the spatial frequency of the stimuli separately for each experimental condition (*valid, invalid*, and *baseline*) and for each cardinal location (left, right, upper, lower) with a Bayesian adaptive psychometric staircase procedure ^70^ used in previous studies ^21–23^. There were a total of 48 independent adaptive procedures (4 per each condition-location combination) for each participant, targeting 75% orientation discrimination accuracy, and each procedure contained 40 trials (**Figure 1B**). To derive the final spatial frequency thresholds, we averaged the estimated thresholds for the 4 staircases. We excluded outlier staircases (1.39 % of all procedures) for which the estimated threshold deviated more than 0.1 log_10_ spatial frequency units from the mean of the other staircases for each condition-location combination. Results showed the same pattern when including the outlier staircases.

To ensure that trials were effectively distributed across the dynamic range and the asymptotes of the psychometric function for the adaptive procedures, and also for the additional fitting analysis (see Materials and Methods-Psychometric function fitting), we calibrated the starting point of the adaptive procedures for the horizontal and vertical meridian based on the initial spatial frequency thresholds measured in the first fixation block (see Materials and Methods-Experimental design). For the horizontal (vertical) meridian, the lowest threshold limit for the adaptive procedure was set to be 0.5 log_10_ spatial frequency units lower than the averaged initial thresholds for left and right (upper and lower) locations, which results in different starting point values (i.e., having a lower threshold limit results in a lower starting value). To account for the pronounced acuity differences between horizontal and vertical meridian observed previously during fixation ^11^, we averaged the spatial frequency thresholds across the left-right and upper-lower locations. Note that the HVA and VMA were evident in the initial spatial frequency threshold measures from the first block, in which we had the same starting point for all locations (mean ±SEM in cpd: left = 9.458 ±0.014, right = 9.609 ±0.013, upper = 6.883 ±0.016, lower = 7.817 ±0.010). Furthermore, the experimental conditions we compared at each location (*valid, invalid, baseline*) all had the same starting values for the adaptive procedures. Thus, the presaccadic effects reported here are independent of the starting values for the adaptive procedures.

### Psychometric function fitting

We fit a psychometric function for proportion correct as a function of spatial frequency (grouped into 20 log-spaced bins), of which the range was flipped in log-space to reflect the increasing psychometric function (**Figure 1C**). For each participant and each location-condition combination, a logistic function was fit to the data using maximum likelihood estimation using the fmincon function in MATLAB’s Optimization Toolbox. For this analysis in particular, we used an offline eye movement data analysis (see Materials and Methods-Eye movement data analysis) because the fitting was conducted after all data had been collected.

### Eye movement data analysis

In addition to the online detection of saccades during the adaptive staircase procedures (see Materials and Methods-Experimental design), we also performed an offline eye movement analysis using an established algorithm for saccade detection ^42^. Saccades were detected based on their velocity distribution using a moving average over 20 subsequent eye position samples. Saccade onsets and offsets were detected based on when the velocity exceeded or fell below the median of the moving average by three standard deviations for at least 20ms. This offline analysis taking into consideration the whole distribution of eye position samples and velocity, and allows for a more detailed evaluation of both fixation and saccadic eye movements. On average 92.6% of trials included in the staircase data also survived the offline analysis (mean ±SEM: 632.917 ±2.745 for *baseline;* 1145.083 ±36.283 for *valid* and *invalid* combined).

### Statistical analysis

The statistical results are based on permutation testing over 1000 iterations with shuffled data. *P* value is the proportion of a metric (*F* score and *t* score) in the permuted null distribution greater than or equal to the metric computed using intact data. Note that the minimum *p* value achievable with this procedure is 0.001. *P* values were FDR corrected for multiple comparisons, when applicable ^71^. The error bars depicting ±1 SEM were Cousineau corrected where indicated ^72^. We also performed Bayesian statistics using the “BayesFactor” package in R (version 0.9.12-4.4). The Bayes factor (BF10) for main effects and interaction effects in the repeated measures ANOVA design were computed by comparing the full model (H1) against the restricted model (H0) in which the effect of interest was excluded from the full model ^73^. BF10 smaller than 1 is in support for the absence of an effect; values 1-3, 3-10, 10-30, 30-100, and >100 indicate anecdotal, moderate, strong, very strong, and extreme evidence for the presence of an effect ^74,75^.

## Additional information and files

### Author Contributions

Conceptualization and methodology: YK, NMH, MC; Software: YK; Formal analysis: YK; Investigation: YK, NMH; Visualization: YK, NMH, MC; Writing – original draft: YK; Writing – review & editing: YK, NMH, MC; Funding acquisition: NMH, MC.

### Data availability

Raw data will be available on the Open Science Framework database (https://osf.io/m8ysa) upon publication.

## Acknowledgements

We thank Marc Himmelberg, Shutian Xue, and Hsing-Hao Lee, as well as other members of the Carrasco lab, for useful comments on the manuscript. This research was supported US NIH National Eye Institute R01-EY027401 to MC and a Marie Skłodowska-Curie individual fellowship by the European Commission (898520) to NMH.

